# Genome-wide profiling of transcribed enhancers during macrophage activation

**DOI:** 10.1101/163519

**Authors:** Elena Denisenko, Reto Guler, Musa Mhlanga, Harukazu Suzuki, Frank Brombacher, Sebastian Schmeier

## Abstract

Macrophages are sentinel cells essential for tissue homeostasis and host defence. Owing to their plasticity, macrophages acquire a range of functional phenotypes in response to microenvironmental stimuli, of which M(IFN-γ) and M(IL-4/IL-13) are well-known for their opposing pro- and anti-inflammatory roles. Enhancers have emerged as regulatory DNA elements crucial for transcriptional activation of gene expression. Using cap analysis of gene expression and epigenetic data, we identify on large-scale transcribed enhancers in mouse macrophages, their time kinetics and target protein-coding genes. We observe an increase in target gene expression, concomitant with increasing numbers of associated enhancers and find that genes associated to many enhancers show a shift towards stronger enrichment for macrophage-specific biological processes. We infer enhancers that drive transcriptional responses of genes upon M(IFN-γ) and M(IL-4/IL-13) macrophage activation and demonstrate stimuli-specificity of regulatory associations. Finally, we show that enhancer regions are enriched for binding sites of inflammation-related transcription factors, suggesting a link between stimuli response and enhancer transcriptional control. Our study provides new insights into genome-wide enhancer-mediated transcriptional control of macrophage genes, including those implicated in macrophage activation, and offers a detailed genome-wide catalogue to further elucidate enhancer regulation in macrophages.

## Introduction

Macrophages are innate immune system sentinel cells that mediate homeostatic and protective functions, including host defence against invading pathogens^1^. Macrophages respond to a wide range of external stimuli by acquiring heterogeneous activation states that exert functional programs tailored for specific microenvironments^2^. A spectrum of macrophage phenotypes has been observed, with macrophages activated in response to interferon-γ, M(IFN-γ), and interleukin-4/interleukin-13, M(IL-4/IL-13), representing two extreme states^3,4^.

M(IFN-γ), often referred to as classically activated macrophages, are pro-inflammatory macrophages characterized by efficient antigen presentation, high bactericidal activity and production of pro-inflammatory cytokines, reactive oxygen and nitrogen intermediates^5,6^. M(IL-4/IL-13), often classified as alternatively activated macrophages, are predominantly regulatory macrophages involved in homeostasis, angiogenesis, wound healing, tissue remodelling and parasitic and bacterial infection^1,2,7-9^. M(IL-4/IL-13) macrophages release anti-inflammatory cytokines and show less efficient antigen presentation and decreased production of pro-inflammatory cytokines^2,7^. Macrophage activation is driven by specific transcriptional changes and is controlled by complex cellular mechanisms^6,10^.

Imbalance in populations of macrophages with opposing pro- and anti-inflammatory roles has been implicated in disease progression^1^. Intracellular pathogen *Mycobacterium tuberculosis*, the causative agent of tuberculosis, interferes with classical activation of macrophages to avoid its anti-bacterial action, and promotes alternative activation state^11,12^. Tumour microenvironments promote phenotypic switches from pro- to anti-inflammatory macrophages, which might contribute to the tumour progression by inhibiting immune responses to tumour antigens^1,2^. Conversely, the phenotypic switch from anti- to pro-inflammatory population of macrophages might contribute to obesity and metabolic syndrome^1,2,13^. Therefore, the development of techniques for manipulation and specific targeting of macrophage populations could ultimately improve diagnosis and treatment of inflammatory diseases^2^. To advance this area of research, the cellular mechanisms responsible for macrophage activation need to be further deciphered.

Gene expression in eukaryotic cells is a complex process guided by a multitude of mechanisms^14^. Precise regulation is required to ensure dynamic control of tissue-specific gene expression and to fine tune the responses to external stimuli^15^. One such level of control is facilitated via regulation of RNA transcription. This process is mediated by a complex transcriptional machinery with its components recognising specific regulatory regions of DNA. Promoters represent a better-characterized class of such regions from which RNA transcription is initiated^16,17^. They act in concert with other cis-regulatory DNA elements, including enhancers, which are believed to play key roles in transcriptional regulation^18^.

Enhancers are defined as regulatory DNA regions that activate transcription of target genes in a distance- and orientation-independent manner^18^. According to the dominant model, transcriptional regulation by enhancers is exerted via direct physical interaction between enhancer and target gene promoter mediated by DNA looping^18,19^. Recent identification of distinct properties of enhancer regions enabled novel approaches to enhancer profiling^18^. Enhancer regions are often distinguished by a specific combination of chromatin marks present at these locations, such as H3K4me1 and H3K27ac^20,21^. Enhancer sequences contain transcription factor binding sites (TFBS) that recruit transcription factors (TFs) to regulate target genes^22,23^. In addition, enhancers are frequently bound by proteins such as histone acetyltransferase p300 and insulator-binding protein CTCF^21,24-26^. Large-scale profiling of these enhancer-associated signatures by chromatin immunoprecipitation followed by sequencing (ChIP-seq)^26,27^ has greatly advanced enhancer identification and enabled systematic and genome-wide enhancer mapping^28,29^. Another group of methods such as chromosome conformation capture (3C)^30^ and its variant Hi-C^31^ has been employed to profile physical DNA contacts, including those between promoters and enhancers^32,33^. However, none of these methods has become a gold standard of enhancer detection, and the field is still actively developing.

Recent studies have led to the unexpected finding that most active enhancers recruit RNA polymerase II and are bi-directionally and divergently transcribed to produce RNA transcripts, referred to as eRNAs^34,35^. While the functionality of eRNA remains controversial, a recent study by Hon et al. showed that many enhancers are transcribed into potentially functional long-noncoding RNAs (lncRNAs) playing a role in inflammation and immunity^36,37^. Recently, quantification of eRNA transcription laid the foundation for a novel method of large-scale enhancer profiling^38^. In their study utilizing cap analysis of gene expression (CAGE)^39^, Andersson et al. performed genome-wide mapping of transcriptional events followed by identification of enhancers based on co-occurrence of closely located divergent transcripts representing eRNAs^38^. The capacity of CAGE to simultaneously profile the expression of eRNAs and genes became an additional advantage, since eRNA production was shown to positively correlate with the production of mRNAs of target genes^34,40^.

These and other studies unravelled the fundamental importance of enhancer regions as DNA regulatory elements in multiple cell types, including macrophages^28,29,35,40,41^. Enhancers are extremely widespread, with an estimation of up to one million enhancers in mammalian genomes^20,23,24,42^. They are major determinants of gene expression programs required for establishing cell type specificity and mediating response to extracellular signals^23,43,44^. Our current understanding of these elements, however, remains incomplete. High tissue-specificity of enhancers is a major hurdle towards establishing a comprehensive catalogue of the full enhancer population^23,43^. Moreover, emerging evidence indicate that enhancers selectively act in a stimuli- or condition-specific manner^45,46^. A major challenge is, therefore, to catalogue enhancers active in different tissues and conditions and link them to target genes.

Recently, we investigated the transcriptional regulatory dynamics of protein-coding and lncRNA genes during M(IFN-γ) and M(IL-4/IL-13) macrophage polarization using CAGE data^10^. We showed that particular TFs, such as Nfκb1, Rel, Rela, Irf1 and Irf2, drive macrophage polarization and are commonly activated but have distinct dynamics in M(IFN-γ) and M(IL-4/IL-13) macrophages^10^. Here, we extended the former study to understand the regulatory influence of enhancers in the macrophage activation process. Our genome-wide *in silico* study aimed at characterizing the transcribed enhancer landscape in mouse macrophages and studying its dynamic changes during M(IFN-γ) and M(IL-4/IL-13) activation. We used CAGE data and enhancer-associated chromatin signature to identify transcribed enhancer regions. We inferred regulatory associations between enhancers and target protein-coding genes using their spatial organisation in topologically associating domains (TADs)^47^ and correlation of CAGE-derived expression in our time-course. With these data, we established a catalogue of transcribed enhancer regions linked to their target genes. This catalogue provides insights into genome-wide enhancer-mediated regulation of transcription in mouse macrophages. Furthermore, we highlight the role of enhancers during macrophage activation and report enhancers driving expression dynamics of known macrophage activation marker genes.

## Materials and Methods

### CAGE data and processing

Mouse genomic coordinates (mm10) and tag counts of CAGE transcription start sites (TSSs) were obtained from the FANTOM5 project^17^ data repository (http://fantom.gsc.riken.jp/5/datafiles/reprocessed/mm10_v2/basic/). Data for 969 mouse samples classified as “primary_cell”, “timecourse”, “tissue”, and “cell_line” were used. The set included 184 macrophage samples profiled by us as described elsewhere^10^, which we used here to construct a macrophage enhancer-promoter interactome (see Supplementary Table S1 for the list of macrophage samples).

The DPI program (https://github.com/hkawaji/dpi1/) was used as described in Forrest et al.^17^ to cluster CAGE TSSs into CAGE peaks. Briefly, the algorithm uses independent component analysis to decompose regions with continuous CAGE signals into separate peaks based on their profile across different samples and tissues. With the default parameters, similarly to Forrest et al.^17^, we obtained a list of all CAGE peaks and a subset of CAGE peaks enriched for promoter-associated signals. The latter file represents a subset of peaks meeting the FANTOM5 ‘robust’ criteria, with a single TSS supported by 11 or more observations and one or more tag per million (TPM) in at least one experiment^17^. These two peak sets were used for identification of enhancers and annotation of protein-coding gene promoters, respectively. Tag counts of all TSSs composing a CAGE peak were summed up to derive a total tag count for that CAGE peak.

### Annotation of protein-coding gene promoters

The set of ‘robust’ CAGE peaks derived by DPI (see above) was used to annotate promoters of protein-coding genes. Ensembl gene models^48^ version 75 downloaded from the UCSC Table Browser^49^ on 11 Aug 2016 was used to obtain coordinates of protein-coding transcripts and genes. We assigned a CAGE peak to an Ensembl protein-coding transcript if its 5’ end was mapped within 500bp of the 5’ end of the transcript on the same strand. The transcript annotation was extended to gene annotation by combining the CAGE peaks associated to all of the gene’s transcripts.

### Calculation of gene and promoter expression

TMM-normalization^50^ of promoter tag counts was performed to derive normalized expression values in a form of tags per million (TPM). We excluded lowly expressed promoters from the analysis and retained only those with expression of at least one TPM in 10% of the macrophage samples. Expression of each gene was derived as a sum of expression of the gene’s promoters. The resulting set included 24,449 promoters of 10,767 protein-coding genes.

### Identification of mouse enhancers with CAGE data

The full set of 3,188,801 DPI-derived CAGE peaks was used for identification of mouse enhancers. CAGE peaks located within 500bp of protein-coding transcript start sites or within 200bp of exons were excluded based on the Ensembl gene models^48^ version 75. This filtering resulted in 1,890,465 CAGE peaks. Next, we used the same strategy as Andersson et al. to infer enhancer regions as clusters of closely located bi-directional divergent CAGE peaks and to derive the corresponding tag counts^38^. The resulting 42,470 regions were deemed mouse enhancer regions. The counts were normalized to tags per million (TPM) using TMM-normalization procedure^50^. Enhancers with non-zero expression in at least 10% of our macrophage samples were deemed transcribed in our macrophage samples.

### Selection of enhancers regulating protein-coding genes in macrophages

Enhancer-specific chromatin signatures were based on ChIP-seq profiling of H3K4me1 and H3K27ac histone marks and were obtained from a study by Ostuni et al.^51^. Transcribed enhancers with at least 1bp overlap with the regions inferred by Ostuni et al.^51^ were retained. Genomic coordinates of TADs in mouse embryonic stem cells were obtained from a study by Dixon et al.^47^. We selected pairs of enhancers and promoters where both features were entirely located within the same TAD. For each of these pairs, we calculated Spearman’s correlation coefficient between expression levels of enhancer eRNA and promoter across our macrophage samples and selected only pairs with positive correlation coefficient and FDR < 10^−4^ (Benjamini-Hochberg^52^ procedure). We considered an enhancer to regulate a gene if it was associated to at least one of the gene’s promoters. All mm9 genomic coordinates were converted to mm10 using the liftOver program (https://genome.ucsc.edu/cgi-bin/hgLiftOver).

### Gene set enrichment analysis

KEGG pathway maps^53^ or GO biological process ontology^54^ were used as sets of biological terms for GSEA. GO terms and associated genes were retrieved using the R package GO.db (Carlson, M. (2015) GO.db: A set of annotation maps describing the entire Gene Ontology). We used hypergeometric distribution to calculate the probability of obtaining the same or larger overlap between a gene set of interest and each biological term^55^. Derived p-values were corrected for multiple testing using Benjamini-Hochberg procedure^52^. As a background, a set of 22,543 Ensembl protein-coding genes (version 75) was used^48^.

### Identification of macrophage-specific features

Normalized TPM expression data were used to calculate a z-score for each of our 184 macrophage samples for each enhancer and gene by subtracting the mean and dividing by the standard deviation of expression values of the same feature in 744 FANTOM5 non-macrophage mouse samples (Supplementary Table S2), similarly to Yao et al.^56^. Enhancers and genes with z-score > 3 (i.e. expressed more than 3 standard deviations above the mean of the non-macrophage samples) in at least 10% of macrophage samples were deemed macrophage-specific.

### TFBS over-representation analysis

TFBS data for mouse were obtained from ENCODE^57^ and HT-ChIP^58^. Raw sequencing data were mapped to the mm10 genome build for each tissue and cell type separately and peaks were called using MACS2^59^. TFBS summits with FDR < 10^−4^ were retained. We used three different background sets: the whole set of identified mouse enhancers, the subset of enhancers not expressed in macrophages, and a set of random genomic regions located within TADs excluding gaps, repeated sequences, Ensembl coding regions, and mouse enhancers identified here. Gap and repeated sequence regions were obtained from the UCSC Table Browser^49^ on 1 Aug 2016 (‘gap’ and ‘rmsk’ tables of mm10 database). Significantly over-represented TFBS were selected based on empirical p-value < 0.01 from a Monte Carlo analysis of 1,000 trials^60^. We retained only TFBS which showed p-value < 0.01 using all three background sets and non-zero expression of the corresponding TF in our macrophages samples.

### Identification of stimuli-responsive features

We calculated a z-score for each of 16 M(IFN-γ) and 16 M(IL-4/IL-13) macrophage samples for each enhancer and gene by subtracting the mean and dividing by the standard deviation of expression values of the same feature in ten non-stimulated macrophage samples, similarly to the approach for identification of macrophage-specific features. Genes and enhancers with z-score > 3 in more than 25% of the corresponding samples were deemed stimuli-responsive. Of associations between stimuli-responsive enhancers and genes, we sub-selected those with a positive Spearman’s correlation of expression in the corresponding activation state.

### Identification of stimuli-specific enhancers

To identify M(IFN-γ)-specific enhancers, we first selected enhancers which were deemed M(IFN-γ)-responsive, but not M(IL-4/IL-13)-responsive. Second, a z-score for each of 16 M(IFN-γ) samples was calculated using 16 M(IL-4/IL-13) samples as a background. Enhancers with z-score > 3 in more than 25% of M(IFN-γ) samples were deemed M(IFN-γ)-specific. Similar strategy was used to infer M(IL-4/IL-13)-specific enhancers.

All analyses made extensive use of the BEDTools utilities ^61^ and the R software (http://www.R-project.org/).

## Results and Discussion

### Identification of transcribed mouse macrophage enhancers

Active enhancers were shown to be bi-directionally transcribed in mammals^34,35^, and eRNAs profiled by CAGE technology^39^ were used before to reliably infer enhancer regions in human^38^. To identify transcribed enhancers in mouse tissue, we used the FANTOM5 collection of CAGE mouse samples^17^ and a strategy developed before^38^ (see Materials and Methods). This approach yielded 42,470 mouse enhancers, with 17,752 enhancers deemed transcribed in our macrophage samples (Figure 1A, Materials and Methods). To refine this set, we sub-selected 11,216 (63%) transcribed enhancers that carry enhancer-specific chromatin signatures (Figure 1A), as determined by ChIP-seq in mouse macrophages^51^ (Materials and Methods). The remaining 6,536 regions did not overlap enhancer regions inferred by Ostuni et al.^51^ and were excluded from further analysis. Notably, of all mouse enhancers not transcribed in macrophages, only 19% carry macrophage enhancer chromatin signatures, highlighting the specificity of enhancers in mouse tissues.

**Figure 1.**
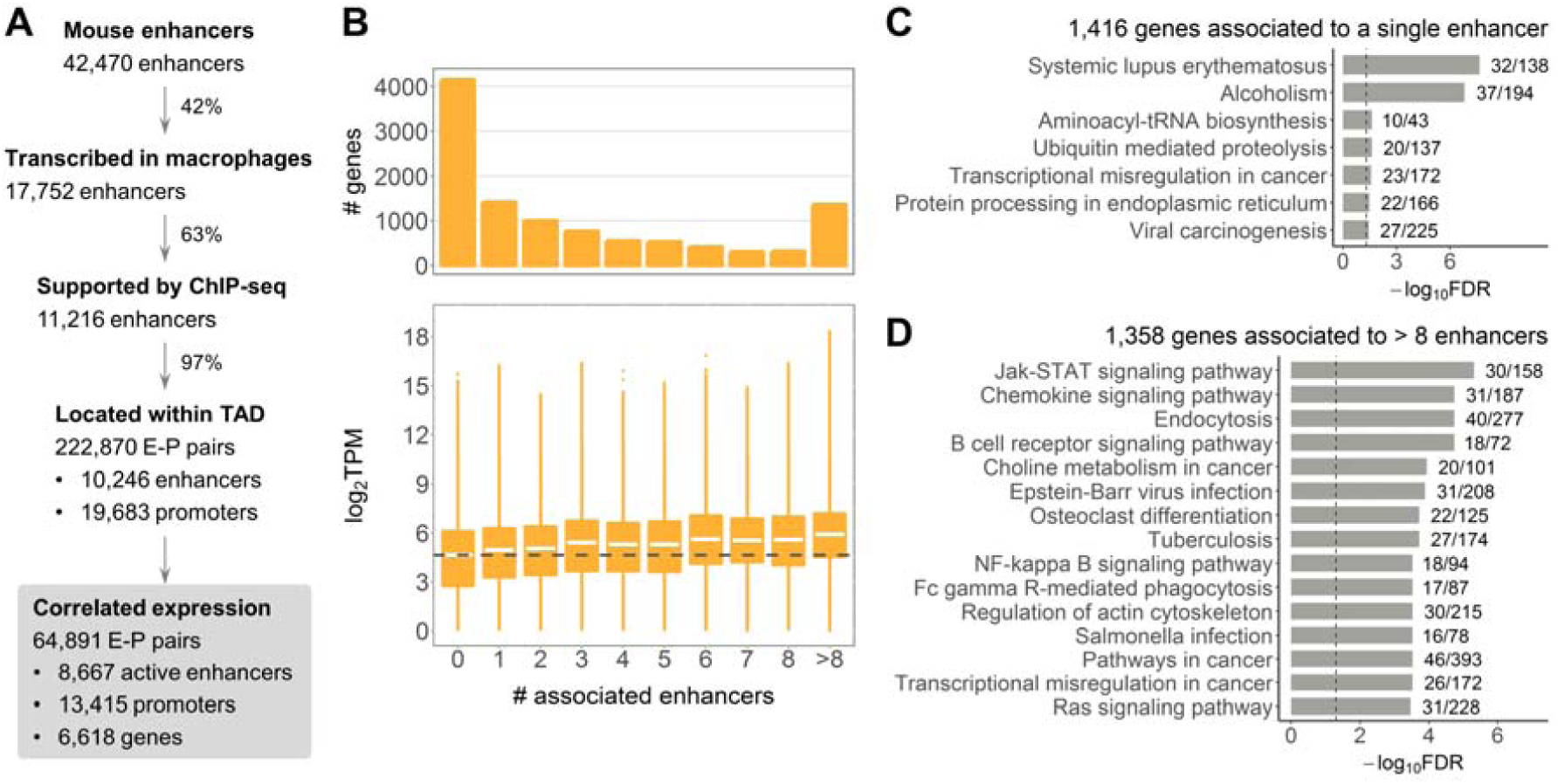
Macrophage enhancer-gene interactome. (**A**) Pipeline for identification of enhancers and enhancer-promoter associations. (**B**) Number and expression of genes associated to different number of enhancers. Dashed line shows median expression of genes not associated to any enhancer. (**C**) KEGG pathway maps significantly enriched for genes associated to a single enhancer, FDR < 0.05. (**D**) Top 15 KEGG pathway maps with the lowest FDR enriched for genes associated to more than 8 enhancers. In (**C**) and (**D**) next to the bars are the numbers of genes in the KEGG pathway covered by our gene list; dashed lines indicate FDR = 0.05.

### Macrophage enhancer-gene interactome

We aimed at studying enhancers that regulate expression of protein-coding genes in macrophages. We first identified pairs of enhancers and promoters located within TADs^47^, since this regulation is thought to be exerted via direct enhancer-promoter contact^18,19^. Thereafter, we refined these pairs using CAGE expression data based on the observation that eRNA and their target expression are positively correlated^34^ (Materials and Methods). This yielded 222,870 TAD-based enhancer-promoter (E-P) pairs, with 64,891 pairs showing significant positive correlation of expression in macrophages (Figure 1A). These correlation-based regulatory associations formed the basis for our further analyses and included 8,667 enhancers deemed active in mouse macrophages. Interestingly, most of the TAD-based E-P pairs showed positive expression correlation (Supplementary Figure S1A), which supports the definition of a TAD as a structural unit favouring internal regulatory interactions^62^. Our filtering approach further selected regulatory associations with the highest correlation (Supplementary Figure S1A), which we considered more reliable. The median distance between enhancers and promoters in the correlation-based E-P pairs was significantly smaller at 191,033nt as compared to 278,735nt for all TAD-based pairs (Supplementary Figure S1B).

We further investigated associations between enhancers and target protein-coding genes (Supplementary Table S3). Of all 10,767 protein-coding genes with CAGE expression (see Materials and Methods), 4,149 genes (38.5%) were not associated to any enhancer in our settings (Figure 1B, upper panel). Given previous evidence of additive action of enhancers^18,63^, we asked whether genes regulated by different numbers of enhancers have different gene expression levels. Genes without associated enhancers were overall lower expressed than genes associated to one (two-sided Wilcoxon signed-rank test p-value < 2.2*10^−16^) or more enhancers. A steady increase in gene expression concomitant with higher numbers of associated enhancers (Figure 1B, Kruskal-Wallis rank sum test p-value < 2.2*10^−16^) was observed, supporting the model of additive enhancer action.

We further asked whether genes associated to different numbers of enhancers within the enhancer-gene interactome show functional differences. Gene set enrichment analysis (GSEA) was performed for gene sets of similar size to avoid a size-related bias (Materials and Methods). The 1,416 genes associated to a single enhancer were enriched for general cellular pathways including “Aminoacyl-tRNA biosynthesis” and “Ubiquitin mediated proteolysis”, as well as a few inflammation-related pathways (Figure 1C). However, in contrast, the 1,358 genes associated to more than eight enhancers showed stronger enrichment for macrophage-related terms, such as “Jak-STAT signaling pathway” and “Chemokine signaling pathway” (Figure 1D). GSEA for 1,306 genes associated to three or four enhancers showed enrichment for a combination of general cellular and macrophage-related pathways (Supplementary Table S4). Finally, the larger set of 4,149 genes not associated to any enhancer showed the strongest enrichment for general cellular pathways (Supplementary Table S5). Hence, a shift towards stronger enrichment for macrophage-related pathways was a concomitant of higher numbers of associated enhancers.

### Macrophage-specific expression

We opted for a similar strategy as Yao et al.^56^ (Materials and Methods) to uncover eRNAs and genes with higher expression in macrophages as compared to other FANTOM5 mouse tissues (further referred to as macrophage-specific). We identified 1,844 macrophage-specific and 8,923 non-macrophage-specific genes (Supplementary Figure S2). These two sets showed significant differences in numbers of associated enhancers, with 65.6% of macrophage-specific genes being associated to more than one enhancer, whereas this proportion dropped to 44.7% for non-macrophage-specific genes (odds ratio 1.99, Fisher’s exact test p-value < 2.2*10^−16^) (Figure 2A). These results were in agreement with our observation of stronger enrichment for macrophage-related functions in genes associated to many enhancers. Similar to the trend observed above, both macrophage-specific and non-macrophage-specific genes showed higher gene expression concomitant with higher numbers of associated enhancers, with non-macrophage-specific genes showing lower expression levels than macrophage-specific ones (Supplementary Figure S3).

**Figure 2.**
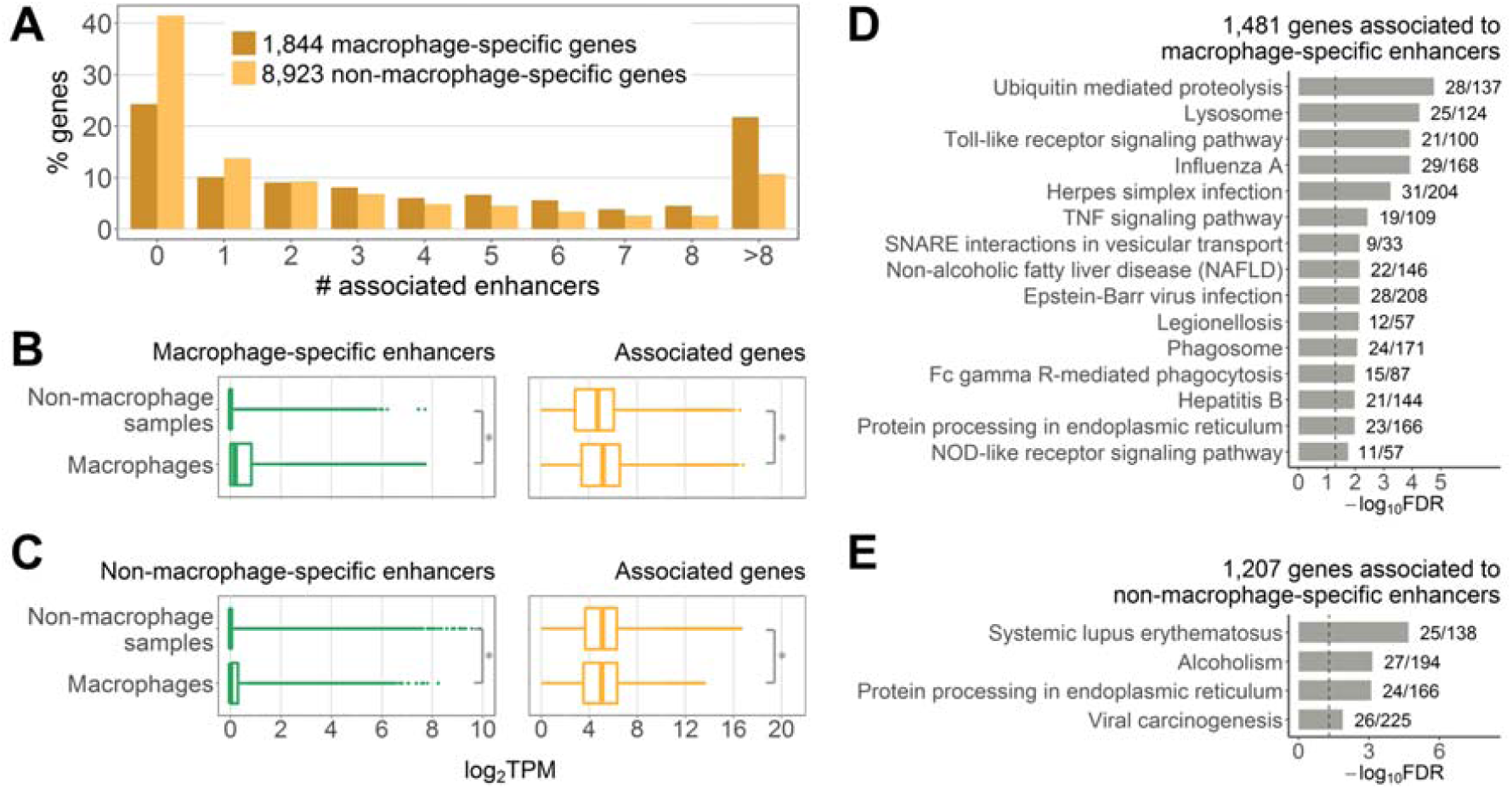
Macrophage-specific enhancer and gene expression. (**A**) Percentage of genes associated to different number of enhancers. (**B**) Expression of 4,739 macrophage-specific enhancer eRNAs and 1,481 associated genes. (**C**) Expression of 3,928 non-macrophage-specific enhancer eRNAs and 1,207 associated genes. In (**B**) and (**C**) expression is shown in 184 macrophage and 774 non-macrophage samples, asterisks denote Wilcoxon rank sum test p-value < 2.2*10^−16^. (**D**) Top 15 KEGG pathway maps significantly enriched for genes associated exclusively to macrophage-specific enhancers. (**E**) KEGG pathway maps enriched for genes associated exclusively to non-macrophage-specific enhancers with FDR < 0.05. In (**D**) and (**E**) next to the bars are the numbers of genes in the KEGG pathway covered by our gene list; dashed lines indicate FDR = 0.05.

Among 8,667 active enhancers, 54.7% were deemed macrophage-specific (Materials and Methods), in agreement with known tissue-specificity of enhancers^23,42,43^. Interestingly, non-macrophage-specific enhancers still showed higher eRNA expression in macrophages as compared to the non-macrophage samples (Figure 2C, left panel). This may be explained by the fact that for this analysis we excluded all enhancers that showed zero eRNA expression in the majority of our macrophage samples (Materials and Methods).

Next, we asked whether these two enhancer sets could regulate genes with different functions. Genes associated exclusively to macrophage-specific enhancers, as well as genes associated exclusively to non-macrophage-specific enhancers were sub-selected. As expected, genes in the former set showed overall higher expression in macrophage samples as compared to the non-macrophage samples (Figure 2B, right panel). In contrast, expression of genes in the latter set was lower in macrophage samples (Figure 2C, right panel). Interestingly, the opposite was observed for non-macrophage-specific enhancers (Figure 2C, left panel). Genes associated to macrophage-specific enhancers were enriched for both general and macrophage-related processes (Figure 2D). This observation reflects the fact that production of macrophage-specific factors and activation of housekeeping processes that facilitate it might be both regulated by the same set of enhancers. Genes associated to non-macrophage-specific enhancers were enriched for only four KEGG pathway maps with FDR < 0.05 (Figure 2E), none of which can be considered a typical macrophage pathway. We obtained consistent results when we repeated the analysis for a subset of 500 genes with the highest expression in macrophages (Supplementary Tables S6-7). Taken together, these findings demonstrate that most of the identified active enhancers in macrophages show macrophage-specific eRNA expression and regulate genes with macrophage-specific as well as general cellular functions.

### Stimuli-induced transcriptional changes

We set out to determine transcriptional changes that were dynamically induced in M(IFN-γ) and M(IL-4/IL-13) mouse macrophages, and to infer enhancers important in these processes (Figure 3A). M(IFN-γ)- and M(IL-4/IL-13)-responsive enhancers and genes were identified as those up-regulated upon stimulation; regulatory associations were retained for pairs with a positive correlation of expression in the corresponding polarization state (Materials and Methods). In this manner, we discovered 115 M(IFN-γ)-responsive enhancers regulating 105 M(IFN-γ)-responsive genes (further referred to as sets E1 and G1), as well as 131 M(IL-4/IL-13)-responsive enhancers regulating 98 M(IL-4/IL-13)-responsive genes (sets E2 and G2) (Figure 3B and Supplementary Tables S8-9). Notably, 77% of E1 and 71% of E2 enhancers were deemed macrophage-specific in our settings. GSEA of G1 and G2 gene sets showed significant enrichment for GO and KEGG terms relevant to immune system and macrophage functions (Figure 3C and Supplementary Figure S4). These results highlight the importance of enhancer regulatory control during macrophage polarization and suggest a striking influence of cytokine stimulation on activation of enhancers, which, in turn, drive some of the transcriptional responses seen during M(IFN-γ) and M(IL-4/IL-13) activation.

**Figure 3.**
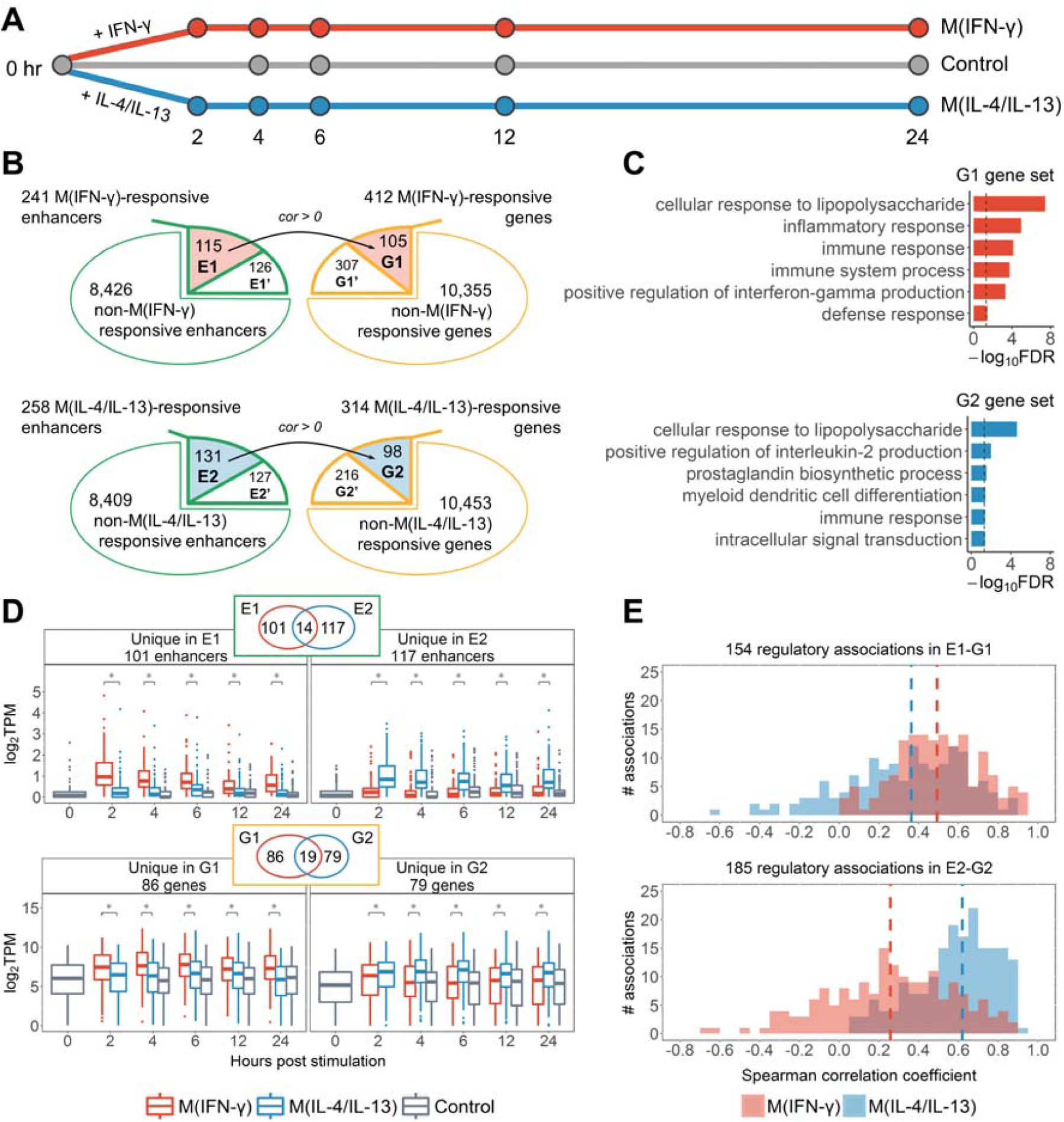
Stimuli-responsive genes and enhancers. (**A**) Time-course data used in this study. (**B**) Enhancer and gene sets. E1 and E2: M(IFN-γ)- and M(IL-4/IL-13)-responsive enhancers regulating M(IFN-γ)- and M(IL-4/IL-13)- responsive genes (G1 and G2), respectively; E1’ and E2’: M(IFN-γ)- and M(IL-4/IL-13)-responsive enhancers regulating non-stimuli-responsive genes; G1’ and G2’: M(IFN-γ)- and M(IL-4/IL-13)-responsive genes not regulated by stimuli-responsive enhancers. Black arrows denote regulatory associations between stimuli-responsive enhancers and genes. (**C**) GO biological process terms enriched for G1 and G2 genes (all terms with FDR < 0.05 for G1; six terms with the lowest FDR for G2 are shown); dashed lines indicate FDR = 0.05. (**D**) Expression of stimuli-responsive enhancer eRNAs (upper panel) and genes (lower panel) unique to M(IFN-γ) and M(IL-4/IL-13). Statistical significance was determined by Wilcoxon signed-rank test, asterisks indicate p-value < 10^−5^. (**E**) Correlation of time-course expression of M(IFN-γ)-responsive (upper panel) and M(IL-4/IL-13)- responsive (lower panel) enhancers and genes. Vertical dashed lines show median values.

M(IFN-γ) and M(IL-4/IL-13) macrophages are known to possess different phenotypes and functions^2^. As expected, G1 and G2 sets had only 19 genes in common. Similarly, a small overlap of only 14 enhancers was observed for E1 and E2 sets. Moreover, enhancers and genes selected as stimuli-responsive for a single activation state showed significant differences in time-course expression in M(IFN-γ) and M(IL-4/IL-13) macrophages (Figure 3D). These data indicate that M(IFN-γ) and M(IL-4/IL-13) macrophages not only differ in their gene expression profiles, but also differ in their active enhancer repertoire that likely drives observed gene expression changes.

Previous studies reported and exploited positive expression correlation of eRNA and target genes^34,38,56^. Hence, we compared expression correlation of E1-G1 and E2-G2 pairs in M(IFN-γ) and M(IL-4/IL-13) macrophages (Figure 3E) to determine how correlations differ between conditions. E1-G1 pairs showed higher correlation in M(IFN-γ) macrophages as compared to M(IL-4/IL-13) (two-sided Wilcoxon signed-rank test p-value = 1.633*10^−6^). Similarly, correlation for E2-G2 pairs was higher in M(IL-4/IL-13) macrophages (two-sided Wilcoxon signed-rank test p-value < 2.2*10^−16^). Such stimuli-specific expression correlation suggests stimuli-specificity of enhancer-gene regulatory associations in macrophages.

### Marker genes of macrophage activation are regulated by stimuli-responsive enhancers

We further asked which known marker genes of classical and alternative macrophage activation^1,2,6,7,64^ were identified in M(IFN-γ) and M(IL-4/IL-13) in our setting (Table 1). Among 20 examined marker genes of classical macrophage activation, we found eight genes in the G1 set; similarly, eight of examined 26 marker genes of alternative activation were found in the G2 set (significant overlap with hypergeometric test p-value < 10^−10^) (Table 1). The G1’ set contained an additional four classical macrophage activation marker genes (Gpr18, Il12b, Il6, Inhba) and the G2’ set an additional three alternative activation marker genes (Il27ra, Klf4, Myc), which, although stimuli-responsive themselves, were not associated to stimuli-responsive enhancers.

**Table 1.**
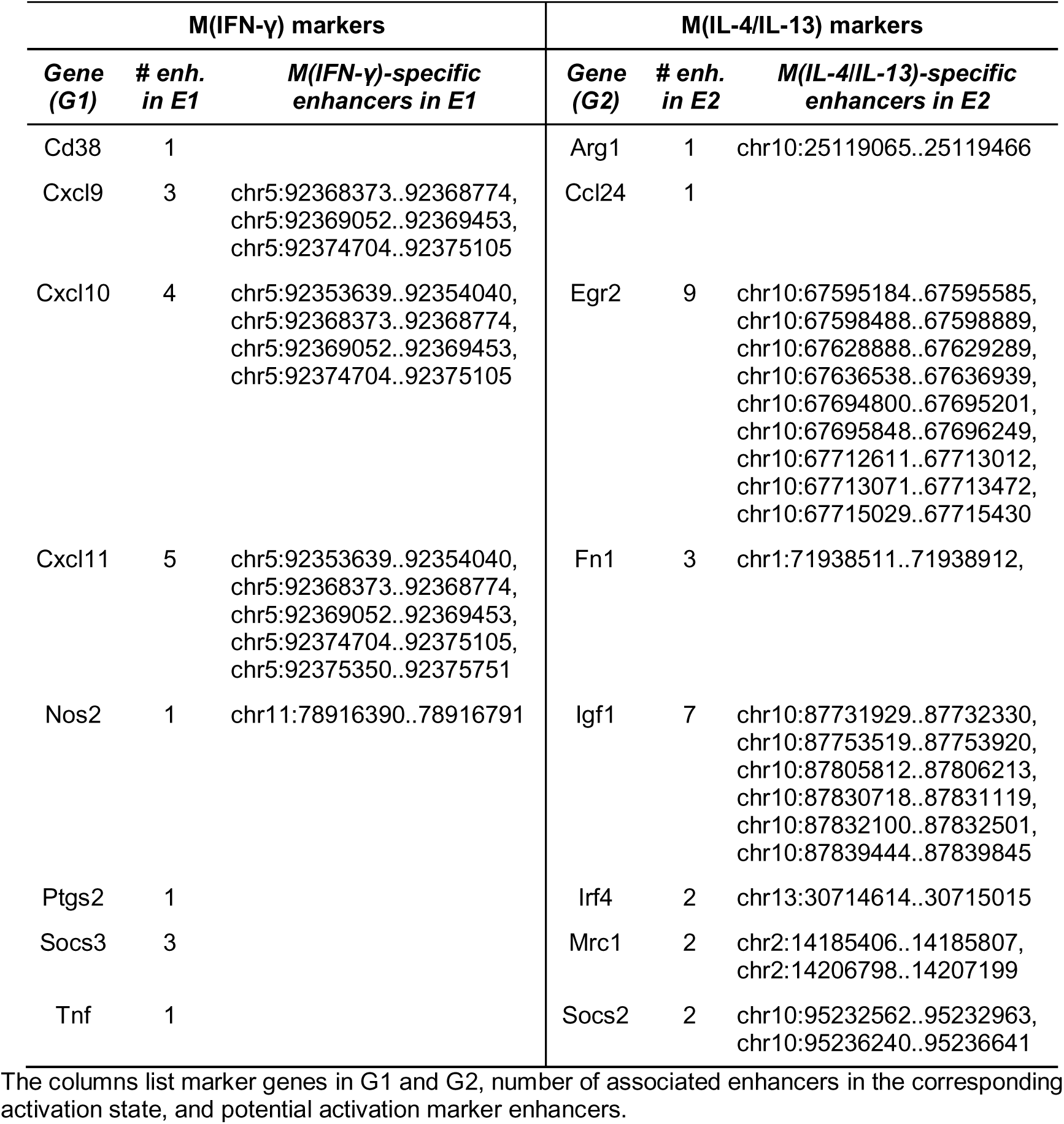
M(IFN-γ) and M(IL-4/IL-13) macrophage activation markers.

| M(IFN-γ) markers | | | M(IL-4/IL-13) markers | | |
| --- | --- | --- | --- | --- | --- |
| Gene (G1) | # enh. in E1 | M(IFN-γ)-specific enhancers in E1 | Gene (G2) | # enh. in E2 | M(IL-4/IL-13)-specific enhancers in E2 |
| Cd38 | 1 |  | Arg1 | 1 | chr10:25119065..25119466 |
| Cxcl9 | 3 | chr5:92368373..92368774,<br>chr5:92369052..92369453,<br>chr5:92374704..92375105 | Ccl24 | 1 |  |
| Cxcl10 | 4 | chr5:92353639..92354040,<br>chr5:92368373..92368774,<br>chr5:92369052..92369453,<br>chr5:92374704..92375105 | Egr2 | 9 | chr10:67595184..67595585,<br>chr10:67598488..67598889,<br>chr10:67628888..67629289,<br>chr10:67636538..67636939,<br>chr10:67694800..67695201,<br>chr10:67695848..67696249,<br>chr10:67712611..67713012,<br>chr10:67713071..67713472,<br>chr10:67715029..67715430 |
| Cxcl11 | 5 | chr5:92353639..92354040,<br>chr5:92368373..92368774,<br>chr5:92369052..92369453,<br>chr5:92374704..92375105,<br>chr5:92375350..92375751 | Fn1 | 3 | chr1:71938511..71938912, |
| Nos2 | 1 | chr11:78916390..78916791 | Igf1 | 7 | chr10:87731929..87732330,<br>chr10:87753519..87753920,<br>chr10:87805812..87806213,<br>chr10:87830718..87831119,<br>chr10:87832100..87832501,<br>chr10:87839444..87839845 |
| Ptgs2 | 1 |  | Irf4 | 2 | chr13:30714614..30715015 |
| Socs3 | 3 |  | Mrc1 | 2 | chr2:14185406..14185807,<br>chr2:14206798..14207199 |
| Tnf | 1 |  | Socs2 | 2 | chr10:95232562..95232963,<br>chr10:95236240..95236641 |

The columns list marker genes in G1 and G2, number of associated enhancers in the corresponding activation state, and potential activation marker enhancers.

Next we inferred potential marker enhancers that regulate marker genes specifically during M(IFN-γ) or M(IL-4/IL-13) activation. Each of the 16 marker genes in G1 and G2 was associated to a minimum of one and maximum of nine enhancers in the E1 and E2 stimuli-responsive sets, respectively (Table 1). Of those, we identified enhancers that were selectively responsive in a single activation state and showed higher expression in this state as compared to the other one (Materials and Methods). A total of 13 M(IFN- γ) and 22 M(IL-4/IL-13) enhancers were inferred as potential activation markers (Table 1).

Interestingly, three marker genes found in M(IFN-γ), Cxcl9, Cxcl10, and Cxcl11 are located within one TAD and are co-regulated by a group of three marker enhancers (Figure 4A-C). These enhancers, along with the two marker enhancers regulating Cxcl10 or Cxcl11 but not Cxcl9 (Table 1) are located in close proximity, in the intronic regions of the Art3 gene (Figure 4C). These enhancer regions were previously reported to show induced RNA polymerase II binding in macrophages upon stimulation with LPS, one of the known classical macrophage activators^35^. In addition, these marker enhancer regions were shown to carry H3K4me1 enhancer histone marks in untreated macrophages^51^. Moreover, H3K27ac modification, associated with active enhancers, is stronger enriched in these regions in M(IFN-γ) as compared to M(IL-4) and untreated macrophages^51^ (Figure 4C), providing further evidence of their functionality in macrophage M(IFN-γ) activation.

**Figure 4.**
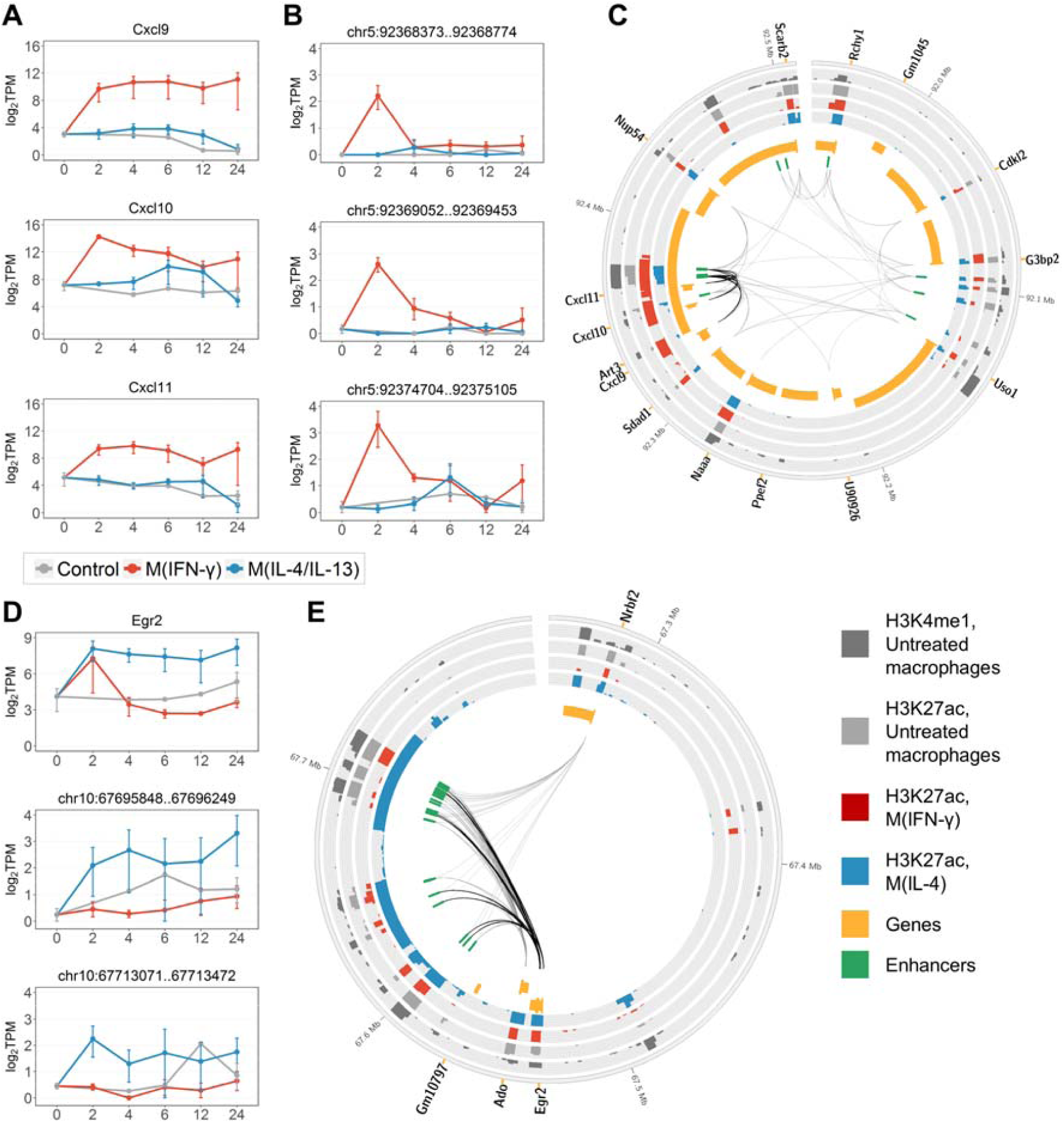
Examples of macrophage activationmarker genes and enhancers. (**A**) Expression of classically activated macrophage marker genes Cxcl9, Cxcl10 and Cxcl11. (**B**) eRNA expression of three potential marker enhancers that co-regulate Cxcl9, Cxcl10 and Cxcl11. (**C**) Genomic region of a TAD containing Cxcl9, Cxcl10, Cxcl11, and associated enhancers. Black links connect the marker genes with the three potential marker enhancers. Grey links denote other enhancer-gene interactions that we identified in macrophages. (**D**) Expression of alternatively activated macrophage marker gene Egr2 and two of M(IL-4/IL-13) marker enhancers associated to Egr2. (**E**) Genomic region of a TAD containing Egr2 and associated enhancers. Black links connect Egr2 with the nine M(IL-4/IL-13) marker enhancers. Grey links denote other enhancer-gene interactions that we identified in macrophages. In (**A**), (**B**) and (**D**) data were averaged over replicates and log-transformed. Error bars are the SEM. In (**C**) and (**E**) genes are split into two tracks based on the strand, wide orange marks denote gene promoters; histone mark tracks show ChIP-seq peaks with the height of -10*log10(p-value), data from^51^.

Among marker genes found in M(IL-4/IL-13), Arg1 as expected, is substantially expressed in M(IL- 4/IL-13)macrophages but has extremely low expression in M(IFN-γ) and untreated macrophages (Supplementary Figure S5A). We found a single M(IL-4/IL-13)-responsive enhancer that might drive expression of Arg1 in M(IL-4/IL-13) macrophages and might serve as a marker enhancer (Table 1 and Supplementary Figure S5B). On the contrary, marker gene Egr2 found in M(IL-4/IL-13), a TF that activates macrophage-specific genes^65^, is associated to as many as nine M(IL-4/IL-13)-specific enhancers (Table 1). Egr2 showed immediate up-regulation in response to both IFN-γ and IL-4/IL-13 stimulation, however, in M(IL-4/IL-13) macrophages the up-regulation sustained for up to 24 hours, whereas in M(IFN-γ) macrophages expression dropped rapidly after 2 hours (Figure 4D, upper panel). Time-course eRNA expression for two Egr2 marker enhancers with the highest expression at 2 and 4 hours is shown in Figure 4D. The distribution of all nine Egr2 marker enhancers within a TAD (Figure 4E) may suggest that the regions identified as nine individual enhancers potentially demarcate fewer regions of stretch enhancers^66,67^. We observed a similar distribution for enhancers of marker gene Igf1, which is known to shape the alternatively activated macrophage phenotype and regulate immune metabolism^68^ (Supplementary Figure S6). Importantly, in both Egr2 and Igf1, marker enhancer regions carried H3K4me1 in untreated macrophages and showed the strongest enrichment with H3K27ac in M(IL-4) as compared to M(IFN-γ) and untreated macrophages^51^ (Figure 4E and Supplementary Figure S6C).

### Transcription factor binding sites are enriched in enhancer regions

To investigate whether our enhancer sets are enriched for known transcription binding sites (TFBS), we performed an over-representation analysis of experimentally determined protein DNA binding sites established through ChIP-seq^57,58^ (Materials and Methods). The sets of macrophage-specific and non-macrophage-specific enhancers are both enriched for binding sites of general factors (p300, Tbp), as well as a range of TFs with well-established roles in macrophages, such as macrophage lineage-determining factor Spi1 (PU.1)^41,69^, Cebpb, required for macrophage activation^70^, and Rela, regulating inflammatory genes^71^ (Supplementary Table S10). Interestingly, TFBS for Spi1 overlap 54.1% of macrophage-specific enhancers, but only 38% of non-macrophage-specific enhancers (overlap ratio of 1.4 for macrophage-specific/non-macrophage-specific enhancers). We observed similar and higher overlap ratios for other functionally important TFs in macrophages, including Stat1, Rela, Irf1, Junb, and Cebpb^6,70^-^72^ (Table 2).

**Table 2.**
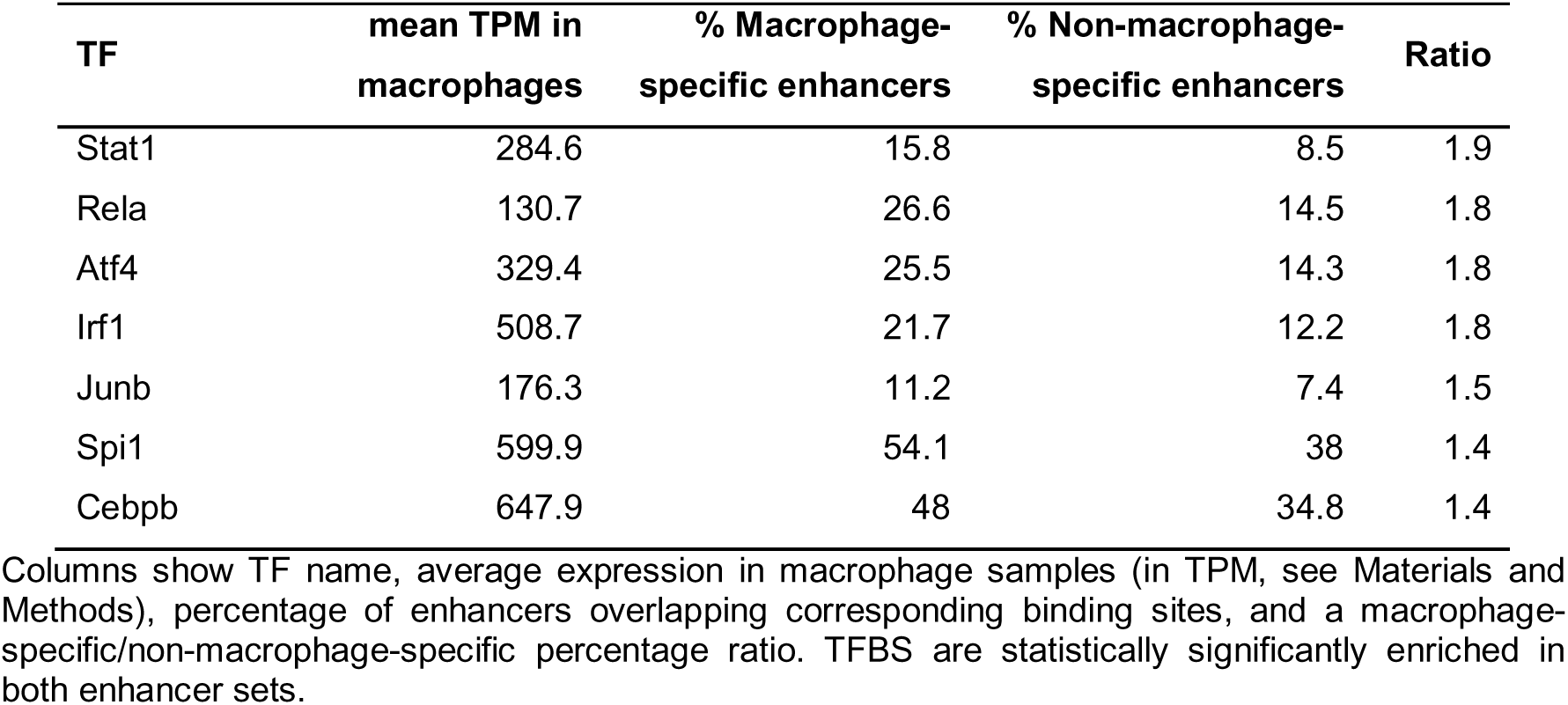
TFs regulating more macrophage-specific than non-macrophage-specific enhancers.

| TF | mean TPM in macrophages | % Macrophage-specific enhancers | % Non-macrophage-specific enhancers | Ratio |
| --- | --- | --- | --- | --- |
| Stat1 | 284.6 | 15.8 | 8.5 | 1.9 |
| Rela | 130.7 | 26.6 | 14.5 | 1.8 |
| Atf4 | 329.4 | 25.5 | 14.3 | 1.8 |
| Irf1 | 508.7 | 21.7 | 12.2 | 1.8 |
| Junb | 176.3 | 11.2 | 7.4 | 1.5 |
| Spi1 | 599.9 | 54.1 | 38 | 1.4 |
| Cebpb | 647.9 | 48 | 34.8 | 1.4 |

Columns show TF name, average expression in macrophage samples (in TPM, see Materials and Methods), percentage of enhancers overlapping corresponding binding sites, and a macrophage-specific/non-macrophage-specific percentage ratio. TFBS are statistically significantly enriched in both enhancer sets.

Similarly, the E1 and E2 stimuli-responsive enhancer sets are enriched for TFBS of known macrophage TFs including Spi1, Cebpb, Rela, and Irf and Stat families^6,73,74^ (Table 3, Supplementary Table S11). Interestingly, TFBS of Stat1, Rela and Irf1, involved in classical macrophage activation^6,75^, overlap a higher percentage of E1 enhancers as compared to E2 and macrophage-specific enhancers (Tables 2-3, and Supplementary Tables S10-11). For instance, Irf1 TFBS overlap 21.7% of macrophage-specific enhancers, 26.7% of E2 but 44.3% of E1 enhancers. In addition, the expression of genes encoding these TFs is higher in M(IFN-γ) as compared to M(IL-4/IL-13) macrophages. Taken together, these results provide an additional layer of support for our regions as functionally important macrophage enhancers and implicate key macrophage TFs in modulating their activity. These findings further reflect that enhancers are selectively activated depending on the transcriptional machinery involved in the cellular response.

**Table 3.** TFs with binding sites enriched in both E1 and E2 enhancer sets.

| TF | mean TPM in<br>M(IFN-γ) | % enhancers<br>in M(IFN-γ) | mean TPM in<br>M(IL-4/IL-13) | % enhancers in<br>M(IL-4/IL-13) | Ratio |
| --- | --- | --- | --- | --- | --- |
| Stat1 | 726.3 | 40.9 | 218.6 | 22.9 | 1.8 |
| Irf1 | 1345.4 | 44.3 | 426.7 | 26.7 | 1.7 |
| Ets1 | 1.4 | 39.1 | 7 | 25.2 | 1.6 |
| Jun | 74.3 | 14.8 | 70 | 9.9 | 1.5 |
| Rela | 131.6 | 42.6 | 102.8 | 29.8 | 1.4 |
| Atf4 | 277.3 | 33.9 | 208.3 | 27.5 | 1.2 |
| Junb | 122 | 13 | 99.7 | 13 | 1.0 |
| Irf4 | 2.3 | 13 | 18.7 | 13.7 | 0.9 |
| Spi1 | 766.3 | 52.2 | 681.7 | 56.5 | 0.9 |
| Cebpb | 424.6 | 51.3 | 262.9 | 59.5 | 0.9 |
| Atf3 | 233.9 | 9.6 | 242.9 | 12.2 | 0.8 |

Columns show TF name, average expression in M(IFN- $\gamma$ ) and M(IL-4/IL-13) macrophages (in TPM, see Materials and Methods), percentage of enhancers overlapping corresponding binding sites, and a M(IFN- $\gamma$ )/M(IL-4/IL-13) percentage ratio.

## Discussion

In this study we investigated the transcribed enhancer landscape in mouse macrophages and its dynamic changes during M(IFN-γ) and M(IL-4/IL-13) activation. Using CAGE data combined with ChIP-seq, we identified 8,667 active enhancers forming 64,891 regulatory associations with protein-coding gene promoters in mouse macrophages. We highlighted tissue- and stimuli-specificity of both enhancers and their regulatory interactions. The enhancer-gene interactome established here supports a model of additive action of enhancers^18,63^, with higher gene expression concomitant with higher numbers of associated enhancers. Moreover, we observed a shift towards stronger enrichment for macrophage-specific biological processes in genes associated to many enhancers. Cytokine stimulation had a striking influence on enhancer transcription, which highlights the importance of enhancers in macrophage activation. In addition, we inferred potential stimuli-specific marker enhancers. Finally, we find that binding sites of inflammatory TFs are enriched in enhancer regions, proposing a link between the response to stimuli and enhancer transcriptional activation.

Several studies have previously reported on enhancer landscape in mouse macrophages. Different populations of tissue macrophages were shown to be highly heterogeneous and to possess distinct sets of active enhancers, as defined by ChIP-seq profiling of histone modifications^28,29^. Studies by Ostuni et al.^51^ and Kaikkonen et al.^40^ used ChIP-seq experiments to uncover enhancers that are established in macrophages *de novo* in response to a range of stimuli. In contrast to previous studies, we combined two complementary data types, transcriptomic (CAGE-derived) and epigenomic (profiled by Ostuni et al.^51^) data, to infer more reliable transcribed active enhancers in mouse macrophages. More importantly, our analysis extended beyond identification of enhancers and characterization of their nearest genes. Here, instead of a widely used linear proximity-based approach^35,38,45^, we employed TAD data to infer enhancer-gene associations. Accumulating evidence suggests that many enhancers regulate distal genes, bypassing the nearest promoter^76,77^. At the same time, TADs have emerged as units of chromatin organisation that favour internal DNA contacts^62^, and the majority of characterized interactions between enhancers and target promoters occur within the same TAD^62,77,78^. Hence, our TAD-based approach enriched with correlation-based filtering enabled us to establish a more reliable mouse macrophage enhancer-gene interactome.

Our interactome covers 8,667 active transcribed enhancers. Of these, 70% overlap RNA polymerase II ChIP-seq peaks in untreated mouse macrophages^51^. Our enhancer regions show significant enrichment for binding sites of histone acetyltransferase p300, an enhancer-associated marker^26^, and known inflammatory TFs. Hence, the regions identified here show a range of known enhancer properties, generally supporting our approach. Most of the active enhancers show macrophage-specific eRNA expression, in line with known tissue-specificity of enhancers^23,42,43^. Kaikkonen et al.^40^ identified mouse macrophage enhancers using ChIP-seq against H3K4me2. Our findings based on CAGE-seq extend their enhancer reportoire by additional 3,974 transcribed enhancer regions. A comparison expanded to non-macrophage enhancers shows that 39.8% of our enhancers overlap a set of *cis*-regulatory elements from 19 non-macrophage mouse tissues identified by Shen et al.^42^. In another recent study, Schoenfelder et al. employed a Capture Hi-C approach to identify enhancers and their target promoters in mouse fetal liver cells and embryonic stem cells^33^. Even though enhancers and enhancer-promoter interactions are known to be highly tissue-specific, their data still support 24.8% of our 64,891 E-P pairs,

Recent reports suggested that genes regulated by multiple enhancers were higher expressed than those regulated by a single enhancer, proposing that enhancers might contribute additively to the expression of their target genes^18,63^. In support of this, we observed a steady increase of gene expression concomitant with increasing numbers of associated enhancers, with the genes not associated to any enhancers showing the lowest overall expression. A study of 12 mouse tissues has reported the enrichment for tissue-specific functions in genes associated to enhancers that transcribe eRNAs as compared to genes associated to non-transcribed enhancers^79^. Jin et al. recently showed that genes that did not interact with distal enhancers were enriched for housekeeping genes and also suggested that cell-specific genes were extensively controlled by *cis*-regulators^80^. We showed in this study that genes associated to many enhancers were more enriched for macrophage-related functions as compared to genes associated to only few or no enhancers. This finding might reflect a more fundamental principle of genome organisation and evolution, such as the importance of multiple enhancers for fine-tuned and redundant control of cell specialization and cell-specific responses.

Studies by Ostuni et al.^51^ and Kaikkonen et al.^40^ revealed stimuli-specific epigenomic changes in enhancer regions in mouse macrophages and introduced a concept of stimuli-specific enhancer activation. In our study, we focused on enhancers and genes that responded to the stimuli with increased expression in order to further investigate this phenomenon. Notably, many stimuli-responsive genes were associated to stimuli-responsive enhancers, highlighting the importance of enhancer regulation in macrophage activation. As expected for such a cell-type-specific process as macrophage activation, most of the responsive enhancers showed macrophage-specific eRNA expression, and genes were enriched for macrophage-specific functions. In addition, our study suggests stimuli-specificity of enhancer-gene regulatory associations in macrophages.

As an important example, we assessed 20 and 26 marker genes of classically and alternatively activated macrophages to characterize their enhancer regulation^1,2,6,7,64^. Of those, seven markers (Ccl20, Fpr2, Ido1; Chi3l3, Chi3l4, Alox12e, Chia) were not expressed in our data. For a total of 16 marker genes, we identified associated enhancers. Moreover, for 11 of them we found enhancers that might regulate these genes specifically in M(IFN-γ) or M(IL-4/IL-13) stimulation (Table 1). Hence, these enhancers present new potential markers for a particular macrophage activation status. Seven additional marker genes, identified as stimuli-responsive, were not associated to any stimuli-responsive enhancer (Gpr18, Il12b, Il6, Inhba in the G1’ set; Il27ra, Klf4, Myc in the G2’ set). The remaining marker genes were not deemed stimuli-responsive. Of those, classically activated macrophage markers Il1b, Cd86, Marco, and Il23a, and alternatively activated macrophage markers Mmp12, Tgm2, Clec4a2, Stab1, F13a1 were associated to at least one enhancer in macrophages. Ccr7, Retnla, Ccl17, Ccl22, Chi3l1, Cxcl13, and Ccl12, were not associated to any enhancers in macrophages.

Moreover, we observed a particular genomic distribution of potential marker enhancers associated to Egr2 and Igf1 marker genes in M(IL-4/IL-13), which suggested that these regulatory DNA regions might represent stretch enhancers. Parker et al., in a recent study, investigated stretch enhancers in human cells and proposed that such extended regions could serve as molecular runways to attract tissue-specific TFs and focus their activity^67^. Similarly to Parker et al., potential stretch enhancer regions identified here were associated to cell-type specific genes and were demarcated by broad H3K27ac signals, specifically higher enriched in M(IL-4) as compared to M(IFN-γ) and untreated macrophages (Figure 4E, Supplementary Figure S6C). Therefore, we propose that stretch enhancers might be involved in the regulation of macrophage activation. However, further studies are required to investigate this phenomenon in more detail.

Our approach inferred M(IFN-γ)- and M(IL-4/IL-13)-responsive enhancers that were strongly enriched for TFBS of known inflammatory TFs. These results are in line with previous reports in mouse macrophages. For example, Spi1 (PU.1) has been extensively studied as a crucial TF involved in macrophage differentiation and transcriptional regulation^41^. Moreover, Spi1 was deemed a pioneering or lineage-determining TF in macrophages, which defines enhancer regions and occupies many enhancers in macrophages^28,35,41,69^. Furthermore, Heinz et al. suggested that collaborative action of Spi1 with Cebpb was required for the deposition of enhancer-associated chromatin marks^69^. Ghisletti et al. reported enrichment for NF-kB (Rel) and Irf TFs in enhancers induced by LPS in mouse macrophages^41^. Likewise, transcribed enhancers induced by LPS and IFN-γ stimulation showed enrichment for NF-kB/Rel, Irf, and Stat1 binding motifs^35^. In addition, we previously showed that TFs including Rela and Irf1 drive expression of protein-coding and lncRNA genes during macrophage activation^10^. Taken together, our results link enhancer activation to the transcriptional program induced by IFN-γ and IL-4/IL-13 stimuli.

In this study, we have established a genome-wide catalogue of enhancers and enhancer-promoter regulatory interactions in mouse macrophages. In contrast to previous studies of enhancer landscape in mouse macrophages, we focused on transcribed enhancers and employed an improved method for identification of enhancer target genes, based on location within a TAD and correlation of expression. Hence, our study represents the most comprehensive analysis of transcribed enhancer activities in mouse macrophages to date and extends current knowledge of transcriptional regulation in macrophages in general and during activation in particular.

## Acknowledgements

This work was supported by grants from the South African National Research Foundation (NRF) and from the Department of Science and Technology, South African Research Chair Initiative (SARCHi) and South Africa Medical Research Council (SAMRC) to F.B.; grant from the Japan Society for the Promotion of Science (JSPS) and National Research Foundation of South Africa to F.B. and H.S.; and grants from the South African National Research Foundation (NRF) Competitive Programme for Unrated Researchers (CSUR) to R.G.

